# Doublet microtubule inner junction protein FAP20 recruits tubulin to the microtubule lattice

**DOI:** 10.1101/2022.11.25.517946

**Authors:** Mamata Bangera, Archita Dungdung, Sujana Prabhu, Minhajuddin Sirajuddin

## Abstract

Motility of organisms involves beating of cilia and flagella that are composed of doublet microtubules. The doublets are made of A- and B-tubules that fuse together at two junctions. Among these, the outer junction is made of tripartite tubulin connections, while the inner junction contains distinct non-tubulin elements. The latter includes Flagellar-associated protein 20 (FAP20) and Parkin co-regulated gene protein (PACRG) that together link the A- and B- tubules at the inner junction. While the structures of doublet microtubules reveal molecular details, their assembly is poorly understood and examining proteins at the junctions can provide important clues. In this study, we purified recombinant FAP20 and characterized its effects on microtubule dynamics by TIRF microscopy. Using *in vitro* reconstitution and cryo-electron microscopy, we conclusively show that FAP20 recruits free tubulin to the existing microtubule, where it mediates flexible lateral interactions between the microtubule lattice and tubulin dimers. Our structure of microtubule:FAP20:tubulin complex partially resembles the inner junction architecture, further providing insights into assembly steps involved in closure of B- tubule in a doublet microtubule.

## Introduction

Biological motility mediated by cilia and flagella beating involves dynein mediated sliding of axonemal doublet microtubules, which are conserved across organisms from unicellular ciliates and flagellates to multicellular eukaryotes (Nicastro *et al,* 2011; Sui & Downing, 2006; Viswanadha *et al,* 2017). Defects in assembly of the doublet microtubule are associated with structural deformities and impairment of ciliary function implicated in many ciliopathies (Fliegauf *et al,* 2007; Oh & Katsanis, 2012). The doublet microtubules are made of a 13-protofilament A-tubule, which resembles cytoplasmic microtubules and an additional, semi-circular B- tubule (with 10 protofilaments) fused to the A-tubule (Satir & Christensen, 2007). Advances in cryo-electron microscopy have led to determination of several high-resolution structures of an intact doublet microtubule from cilia/flagella (Khalifa *et al,* 2020a; Ma *et al,* 2019). These models have revealed intricate details about the doublet microtubule architecture, various microtubule inner proteins decorating both the A- and B-tubule as well as the outer and inner junctions between the A- and B-tubule. The outer junction is entirely mediated by tubulin components that form a tripartite lateral interaction between A- and B- tubule protofilaments (Ichikawa *et al,* 2017; Khalifa *et al,* 2020a; Ma *et al,* 2019). The assembly of outer junction has been demonstrated *in vitro* in presence of excess salts as well as in the absence of tubulin carboxy terminal tails. (Euteneuer & McIntosh, 1980; Schmidt-Cernohorska *et al,* 2019). However, the partially assembled B-tubule in this intermediate does not fuse with the A-tubule and presumably requires proteins of the inner junction to form the complete doublet microtubule (Khalifa *et al*, 2020a; Nicastro *et al*, 2011). The doublet microtubule inner junction had long been suspected to be mediated via non-tubulin subunits (Nicastro *et al*, 2011) and the components were later identified from high-resolution structures (Khalifa *et al,* 2020a; Ma *et al,* 2019). An alternating set of FAP20 and PACRG protein units forms the core of the inner junction (Dymek *et al*, 2019; Khalifa *et al*, 2020a; Ma *et al*, 2019) and recent high-resolution cryo-electron tomography structures reveal additional protein subunits such FAP45, 52, 106, 126 and 276 further strengthening the inner junction (Khalifa *et al*, 2020a).

Mutational studies in *Chlamydomonas* reveal that although PACRG and FAP20 proteins are not essential for the formation of flagella, their null mutants show abnormalities in the structure of axonemes (absence of beak structures and inner dynein arms) and severe defects in motility of the organisms (Dymek *et al,* 2019; Yanagisawa *et al,* 2014). Similar effects are also observed in case of depletion of a homologue of FAP20 in *Paramecium,* known as Bug22 (Laligné *et al,* 2010). Bug22 mutants in Drosophila show abnormalities associated with ciliopathies, neuronal defects and impairment of sperm differentiation (Mendes Maia *et al,* 2014). Depletion studies also propose the link between Bug22 and posttranslational modifications of the axonemal doublet in Drosophila as well as in human RPE1 cells (Mendes Maia *et al,* 2014). Interestingly, the absence of either PACRG or FAP20 causes slower sliding velocities with respect to dynein but the effect on the double mutant is not cumulative (Dymek *et al,* 2019). These observations suggest that PACRG and FAP20 play a role in later stages of the inner junction assembly and contribute towards maintaining the structural integrity and functional aspects of the axoneme. So far, *in vitro* studies of inner junction proteins has been limited to experiments with recombinant PACRG (Ikeda, 2008; Khan *et al,* 2021) and there is little biochemical evidence to show the direct role of FAP20 in doublet formation and maintenance. To address this, we have purified a recombinant version of FAP20 and characterized its biochemical properties towards microtubules using *in vitro* reconstitution studies. Our studies with light and electron microscopy show that FAP20 recruits tubulin to the microtubule lattice independent of GTP. The reconstitution of FAP20 binding to a growing microtubule allowed us to identify flexible interactions in the microtubule:FAP20:tubulin complex and propose a model for inner junction assembly.

## Results

### FAP20 binding stabilises the microtubule lattice

The properties of FAP20 binding to microtubules *in vitro* have not been reported so far and to study this we cloned, expressed and purified a recombinant version of full length FAP20 from *Chlamydomonas reinhardtii* by nickel-histidine affinity chromatography (See Methods for details, Supplementary Figure 1A). First we performed copelleting assays with purified FAP20 and microtubules polymerised from goat brain tubulin in the presence of different nucleotides (Supplementary Figure 1B and 1C). We found that FAP20 binds to all of the microtubules *in vitro*, suggesting that FAP20 is insensitive to minor lattice perturbations due to different nucleotide states (Zhang *et al,* 2015) (Supplementary Figure 1C). To assess whether FAP20 affects microtubule growth and stability we then carried out microtubule growth assays in the presence of FAP20 (See Methods for details, Figure 1A, Supplementary Movie 1). From the kymograph analysis of TIRF images, we found that at lower concentrations, FAP20 did not induce any significant effect on the growth rate and growth length of the microtubules (Figure 1B and 1C, Supplementary Figure 2A and 2B). However, the growth rates and growth lengths of microtubules increased significantly in presence of 2.5 μM and 5μM FAP20. The microtubule growth rates in the absence and presence of 2.5 μM and 5uM FAP20 were 0.54 μm min^-1^,0.89 μm min ^-1^ and 0.88 μm min^-1^ respectively (Figure 1B). The microtubule growth lengths recorded were 2.93 μm in absence of FAP20 and 6.00 μm and 6.32 μm in presence of 2.5μM and 5μM FAP20 respectively (Figure 1C). Another striking observation apparent from the kymographs was the reduced number of catastrophe events as the concentration of FAP20 was increased in the microtubule growth assays (Figure 1A). Quantification of the frequency of catastrophe events showed marked decrease in a concentration dependent manner. In the absence of FAP20, and with 0.05 μM and 0.25μM FAP20, 80.4%, 68.3% and 54.66% of the analysed microtubules exhibited regular catastrophe events respectively (Figure 1D and Supplementary Figure 2C). At higher concentrations of FAP20 (2.5uM and 5uM) almost no catastrophe events could be observed (Figure 1A and 1D). Thus the recombinant FAP20 protein makes stabilising interactions with the microtubule lattice increasing the growth rate, reducing the number of catastrophe events and eventually preserving the growing microtubule.

**Figure 1.**
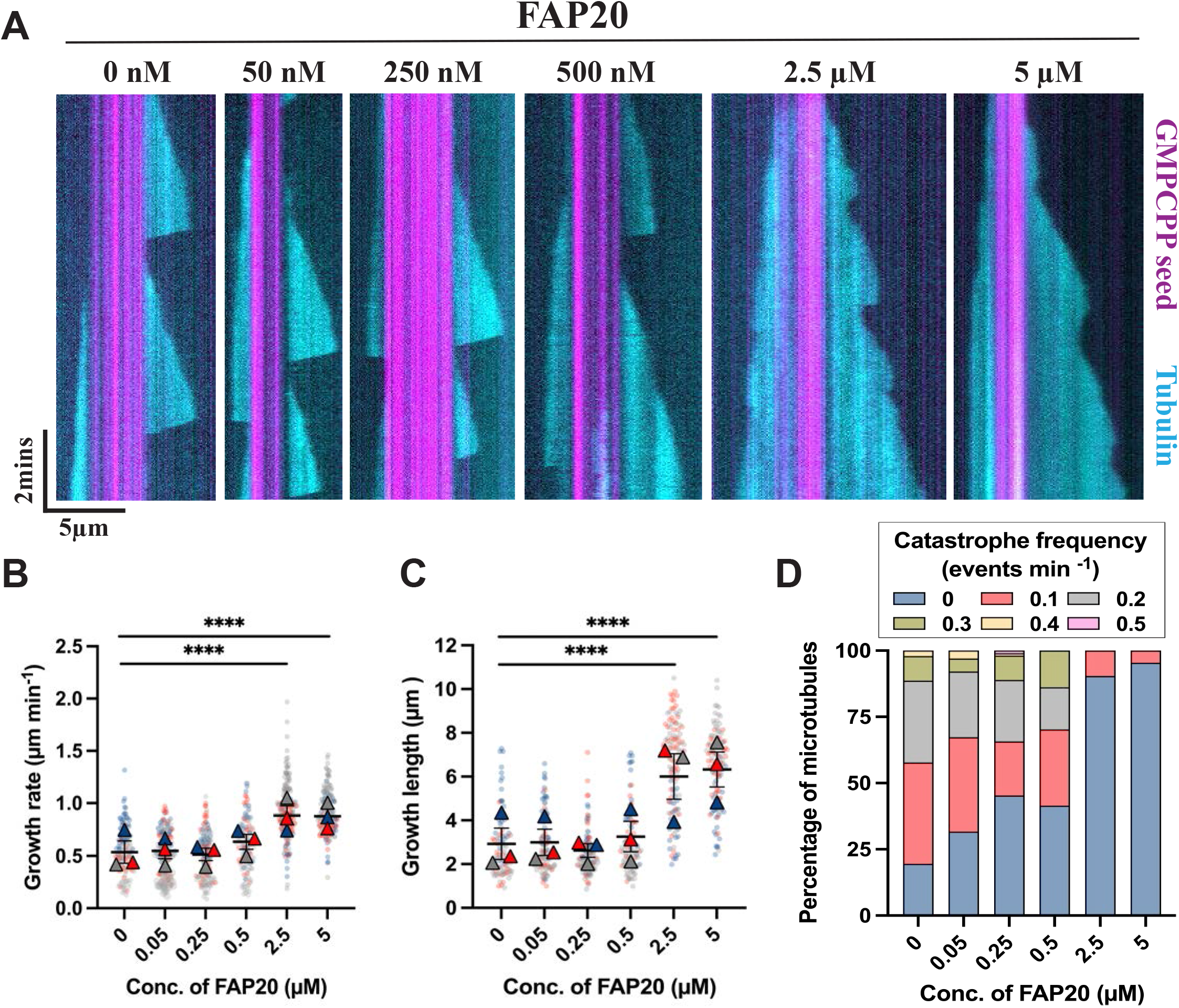
Microtubule dynamics in the presence of FAP20. **(A)** Representative kymographs showing microtubule dynamics in the presence of different concentrations of FAP20. **(B) and (C)** SuperPlots of microtubule plus-end growth rates and growth lengths in the presence of 10μM tubulin and increasing concentrations of FAP20, as indicated. Average values from three independent experiments are shown as coloured triangles and individual values from each experimental set are shown as circles in their respective muted colours. Error bars represent standard error of the mean from three independent experiments. (N = 135: 0μM FAP20, 130: 0.05μM FAP20, 137: 0.25 μM FAP20,118: 0.5μM FAP20, 164: 2.5μM FAP20, 150: 5μM FAP20 for growth rate measurements and N = 97: 0μM FAP20, 101: 0.05μM FAP20, 108: 0.25 μM FAP20, 94: 0.5μM FAP20, 125: 2.5μM FAP20, 110: 5μM FAP20 for growth length measurements. ****p< 0.0001, Kruskal-Wallis test followed by Dunn’s multiple comparisons test) **(D)** Frequency plot showing percentage of microtubules undergoing different catastrophe frequencies at each FAP20 concentration. Data is obtained from three individual experiments (N = 97: 0μM FAP20, 101: 0.05μM FAP20, 108: 0.25 μM FAP20, 94: 0.5μM FAP20, 125: 2.5μM FAP20, 110: 5μM FAP20). For individual values see Supplementary Figure 2 and Source Data 1.

### FAP20 recruits free tubulin to microtubule lattice

In our microtubule growth assays, at higher concentrations of FAP20 we observed fluorescence signal from cy5-labelled free tubulin accumulating at the existing microtubule lattice, both GMMPCP seeds as well as the newly polymerized microtubules (Figure 2A and Supplementary Movie 1). We reasoned that FAP20 promotes lateral association of free tubulin dimers to the microtubule lattice and investigated the role of GTP in this phenomenon. Under the same conditions without GTP, similar cy5-labelled free tubulin accumulation was observed (Figure 2B and Supplementary Movie 2) suggesting that the FAP20 protein is the basic unit that links the microtubule lattice to the soluble-tubulin dimer. Although we observed different patterns of tubulin association with the GMPCPP seeds in presence of GTP (Supplementary Figure 3A, B and C), the pattern of arrangement was found to be regular and continuous in case of tubulin recruitment by FAP20 when no GTP was present (Supplementary Figure 3D and 3E).

**Figure 2.**
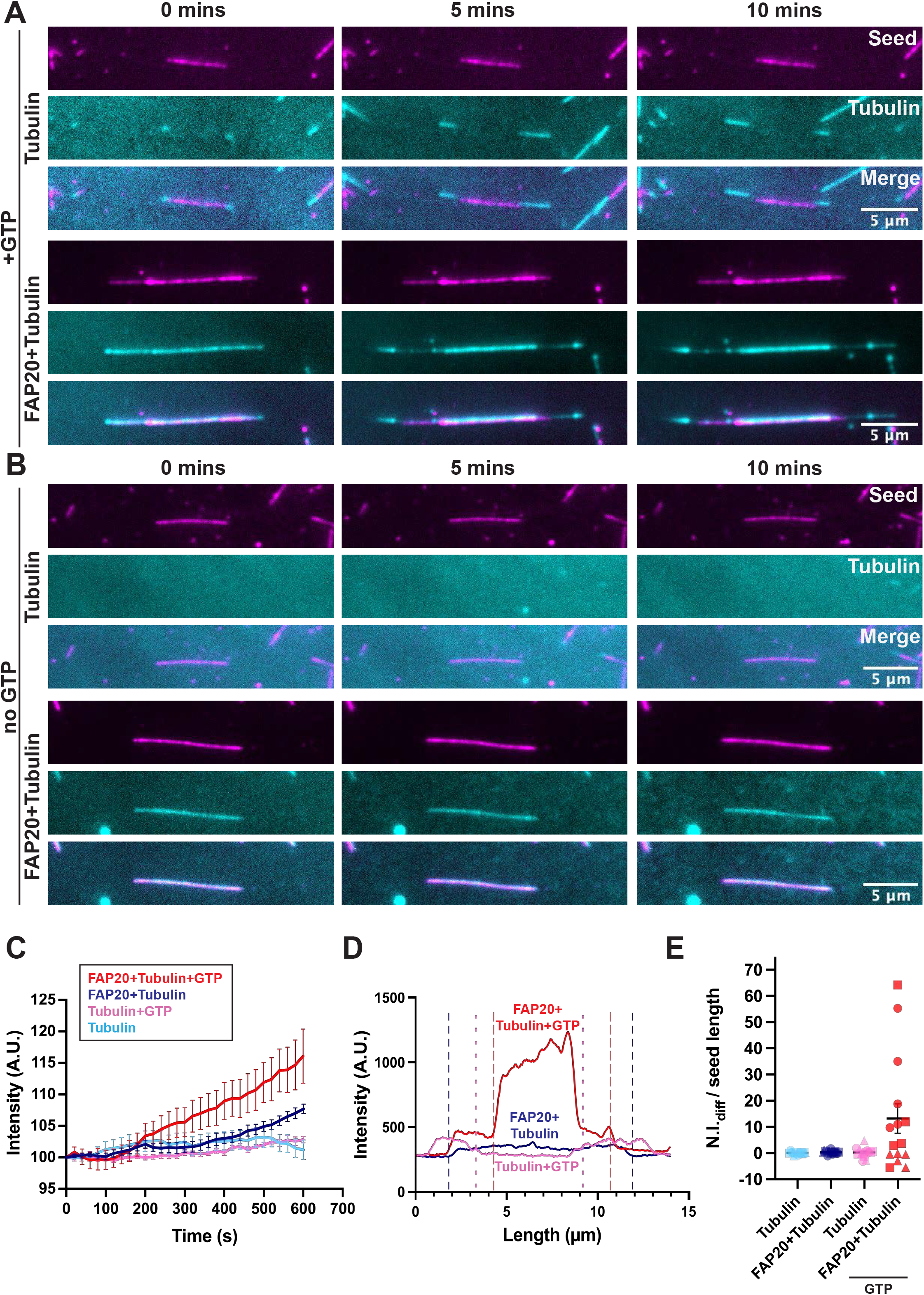
FAP20 mediated recruitment of tubulin to the microtubule lattice. **(A)** Representative TIRF images of GTP microtubules grown from GMPCPP seeds (magenta) in presence of 10μM tubulin (cyan) with and without 5μM FAP20 at different time points, as indicated. **(B)** Representative TIRF images of GMPCPP seeds (magenta) in presence of 10μM tubulin (cyan) with and without 5μM FAP20 in the absence of GTP at different time points, as indicated **(C)** Mean fluorescence intensity of cy5 labelled-free tubulin along the GMPCPP seed with time for 10 minutes in the presence of 10μM tubulin (light blue), +GTP (pink), +FAP20 (dark blue) and +FAP20 with GTP (red). Error bars represent standard error of mean from three independent experiments. **(D)** Fluorescence intensity line scans (excitation at 561nm) along the length of a microtubule in the presence of 10μM tubulin (pink) and GTP, + 5μM FAP20 without GTP (dark blue), + 5μM FAP20 with GTP (red). The dotted lines represent the ends of the GMPCPP seeds for the respective experiments. **(E)** Differences in final and initial normalised intensity per length (μm) of GMPCPP seed are shown in presence of 10μM tubulin (light blue), +GTP (pink), +FAP20 (dark blue) and +FAP20 with GTP (red). Values from each experimental set are represented as different symbols and error bars represent standard error of mean from three independent experiments (N=5 for each, from three independent experiments).

To gain further understanding into the pattern of tubulin recruitment, we measured the fluorescence signal associated with bound tubulin dimers over time (Figure 2C) and the length of the microtubule (Figure 2D). We observed a uniform increase in tubulin recruitment over time on the GMPCPP stabilised seeds, with an increased recruitment in the presence of GTP compared to its absence (Figure 2C). This effect was also evident from intensity analysis over the length of the microtubule, where we observed several fold increase in fluorescence intensity represented by the area under the curve corresponding to the GMPCPP seed only in the presence of GTP (Figure 2D). Quantification of the difference between initial and final normalised intensities of recruited tubulin to the GMPCPP seed showed a large distribution of values (Figure 2E) suggesting no directed pattern of recruitment in presence of GTP. The FAP20 mediated tubulin recruitment occurs rapidly as the reaction is setup and the initial stages could not be captured using TIRF imaging (Figure 2A and 2B). This is particularly evident in absence of GTP, where the difference in normalized fluorescence intensities between initial and final time points of the assay is negligible as compared to in presence of GTP (Figure 2E). The tubulin recruitment saturates almost immediately in absence of GTP, while in its presence the recruited tubulin dimers act as nucleation points for a steady state lateral association. In summary, our TIRF imaging experiments have uncovered a unique property of FAP20, where it mediates binding of a tubulin dimer to the microtubule lattice in a GTP independent manner.

### FAP20 mediates flexible interaction of microtubule lattice with tubulin dimer as well as protofilaments

We next performed cryo-electron microscopy analysis to understand the FAP20 mediated tubulin recruitment on to the microtubule lattice. Samples were prepared by adapting the steps derived from our TIRF assay on Transmission Electron Microscopy grids, which were then imaged in an electron microscope (See Methods for details). In the absence of FAP20, we observed undecorated GMPCPP microtubules with a diameter between 25 – 30 nm (Figure 3A). In the presence of GTP and FAP20,we observed microtubule-like structures with twice the diameter ~50 nm (Figure 3B), indicating tubulin accumulation as observed in the TIRF images (Figure 2A). Analysis of the raw and filtered images as well as the power spectra suggests presence of additional layers of protofilaments associated with each other on a conventional microtubule lattice structure (Supplementary Figure 4A). Attempts to obtain a consensus high resolution 3D reconstruction of this structure failed due to heterogeneity of the sample and the complexity of the arrangement in the higher order structure (Supplementary Figure 4B). However, a low resolution structure could be determined, which shows multiple stacks of protofilaments with a hollow centre similar to the diameter of a conventional microtubule (Supplementary Figure 4B).

**Figure 3.**
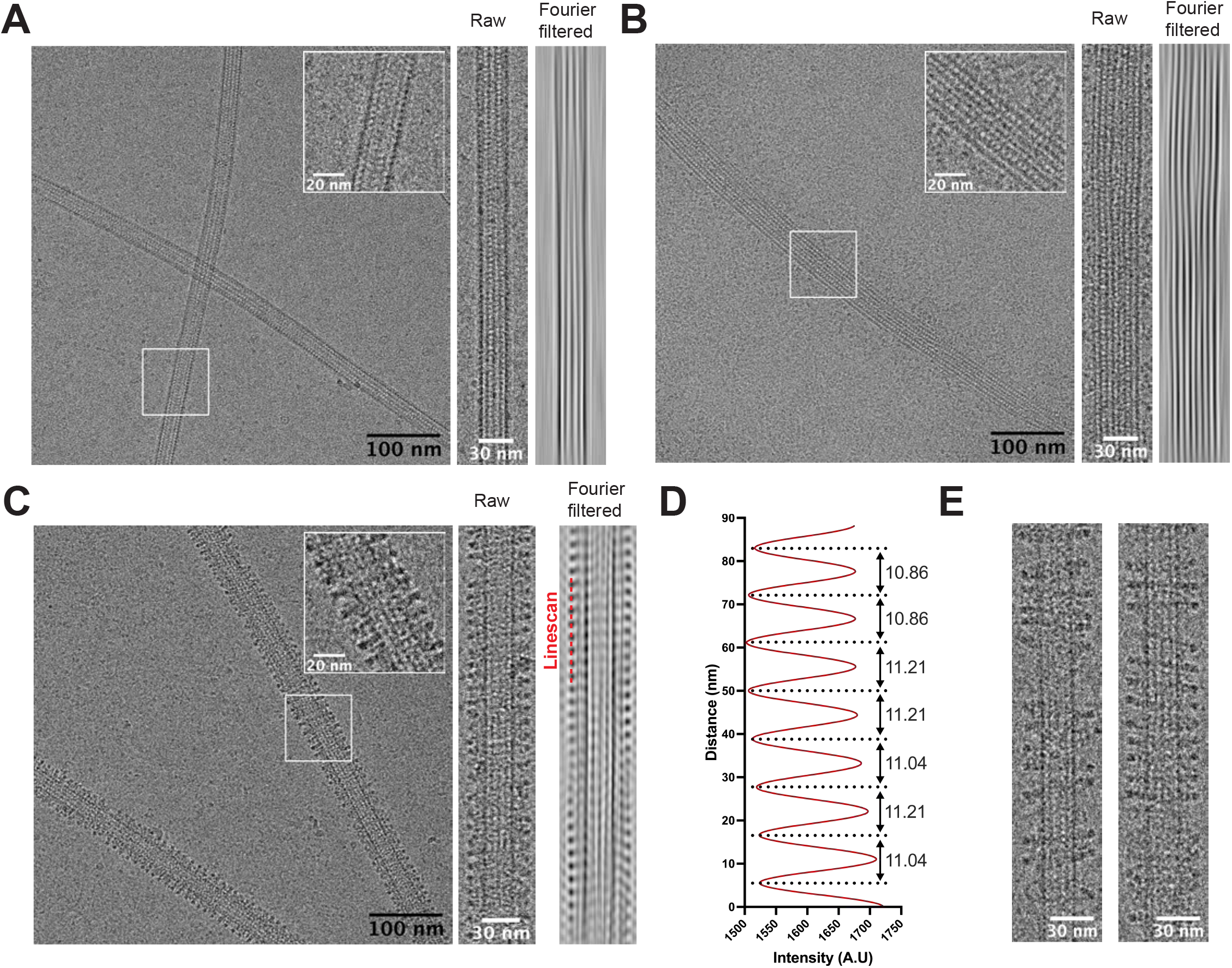
Ultrastructural analysis of FAP20 mediated tubulin recruitment. **(A)-(C)** Representative cryo-electron micrographs showing GMPCPP microtubules (A), +FAP20 and 10μM tubulin with GTP (B) and +FAP20 and 10μM tubulin (C). Inset: Zoomed in version of a section of microtubule outlined by a white box in the micrographs. Right: Longitudinal sections of a single microtubule segment with its Fourier filtered images to highlight the differences. **(D)** Line scan plot of intensity distribution along the decorated region of microtubule+FAP20 and tubulin selected by the red dotted line in (C). **(E)** Straightened segments of micrographs showing heterogenous FAP20+tubulin ring/spiral formation along the microtubule lattice in absence of GTP.

To obtain a more homogenous sample we then turned our attention to FAP20 mediated tubulin recruitment in the absence of GTP. In the absence of GTP, the electron micrographs showed presence of the microtubule lattice (20 – 30 nm) decorated with rings and/or spirals (Figure 3C). From the line scan analysis of the filtered images (Figure 3D) and the layer lines in the power spectra of the helical segments (Supplementary Figure 4A), a 10-11 nm regular interval of rings/spirals structures could be observed. Since the *in vitro* reconstitution system contains only microtubule, free tubulin and FAP20, and the TIRF images show tubulin recruitment on existing microtubule lattice, we deduce that the rings/spirals observed here are higher order structures composed of microtubule, FAP20 and tubulin dimers. The interaction between FAP20, tubulin dimers and microtubule however appears to be flexible as the rings/spirals were found at different angles with respect to the microtubule axis and the vertical spacing between the rings/spirals, depending on the region analysed might actually range around 11 nm (Figure 3E). Hence, we conclude that FAP20 mediates flexible interactions between the microtubule lattice and tubulin dimers in the absence of GTP. This association is retained with protofilaments when GTP is added to the sample.

### Binding of FAP20 to the microtubule is specific

From the TIRF experiments and electron micrographs, the microtubule:FAP20:tubulin structure (in absence of GTP) appeared to be less heterogenous compared to microtubule:FAP20:tubulin:GTP complex (Figure 2E, 3B and 3C). Therefore, we attempted 3D reconstruction of microtubule:FAP20:tubulin complex to deduce its architecture and possible molecular interactions (See Methods for details and Supplementary Figure 5A). A low resolution reconstruction of the complex revealed that it is made up of three concentric layers corresponding to the microtubule (inner), FAP20 (intermediate) and tubulin dimers (outer)(Figure 4A). From initial assessment, the outer and inner concentric layers form 1-start and 3-start helices (Figure 4A), in line with our power-spectra analysis where we observed an additional layer line for microtubule:FAP20:tubulin complex (Supplementary Figure 4A). This posed a challenge during processing as imposing helical symmetry using a single set of helical parameters was not possible. To improve the resolution of the complex, we performed signal subtraction for the different layers and refined them individually with their corresponding sets of helical parameters (Supplementary Figure 5A). This process yielded a 3.9 Å resolution cryoEM map of the microtubule lattice (Figure 4B, Supplementary Figure 5B). However the particles with subtraction focussed on the FAP20 layer and outer tubulin layer did not yield EM maps with defined structures for the intermediate and outer rings/spirals (FAP20 and tubulin dimer unit), possibly because of heterogeneity and flexibility in these layers (Figure 4B, Supplementary Figure 5A and 5B). In particular, the FAP20 layer was very poorly resolved and did not show any discernible features that could aid in fitting the model of FAP20 obtained from cryo-electron microscopy studies on the isolated doublet (Khalifa *et al*, 2020a; Ma *et al*, 2019). The outer layer corresponding to the tubulin dimers showed distinct blobs of density that could accommodate tubulin dimers horizontally (Figure 4B). The horizontal protofilament along the ring/spiral showed better fit into the density compared to the vertical alignment of tubulin dimers along the microtubule axis (Figure 4C). In order to obtain a composite model of the microtubule:FAP20:tubulin interaction, we attempted AlphaFold2 (Evans *et al,* 2021; Jumper *et al,* 2021a) based structure prediction of complex of FAP20 and tubulin dimer and compared the predicted models with structures of the fully assembled doublet determined by cryo-electron microscopy. The AlphaFold2 model of FAP20 bound to a tubulin dimer showed high confidence (pTM+ipTM=0.91) and was similar to the interaction of FAP20 with alpha tubulin from A1 protofilament in the atomic models of the doublet (Supplementary Figure 5C and 5D). The distinct spacing of FAP20 mediated tubulin rings in the structure, high confidence values (pTM+ipTM between 0.89 to 0.91) for top 5 models obtained from Alpha Fold (Supplementary Figure 5D) and buried surface area (865.2 Å^2^) suggest a specific binding mode of FAP20 towards the alpha tubulin in the microtubule lattice (similar to A1 protofilament in the doublet microtubule). Using the above information, we obtained a reasonable fit of this model of tubulin and FAP20 (from AlphaFold2) in the electron density maps of subtracted particles corresponding to the microtubule and FAP20 layers and fit the horizontal tubulin protofilament in the map corresponding to the outer tubulin layer (Figure 4D). Examination of the combined model revealed that the distance between microtubule lattice and tubulin dimer was 6 nm with no linking connections in the electron density map between FAP20 and the microtubule lattice or tubulin dimer, indicating high flexibility of the interactions. The interaction between FAP20 and tubulin dimers of the outer helix does not resemble the FAP20-B-tubule interaction determined from the cryoEM structure of the doublet microtubule (Khalifa *et al*, 2020b; Ma *et al*, 2019). Analysis of the interface of FAP20 with the B-tubule in the doublet structure (PDB ID: 6u42) gave low values of buried surface areas (FAP20: alpha tubulin-135.2 Å^2^, FAP20: beta tubulin-262.4 Å^2^) suggesting weak interactions possibly stabilised by other proteins of the inner junction. Additionally the map corresponding to the FAP20 layer showed residual gaps after fitting of the atomic models of FAP20, which can incorporate a slightly extended version of FAP20 or another molecule of FAP20 and hence might represent an intermediate in the formation of the inner junction.

**Figure 4.**
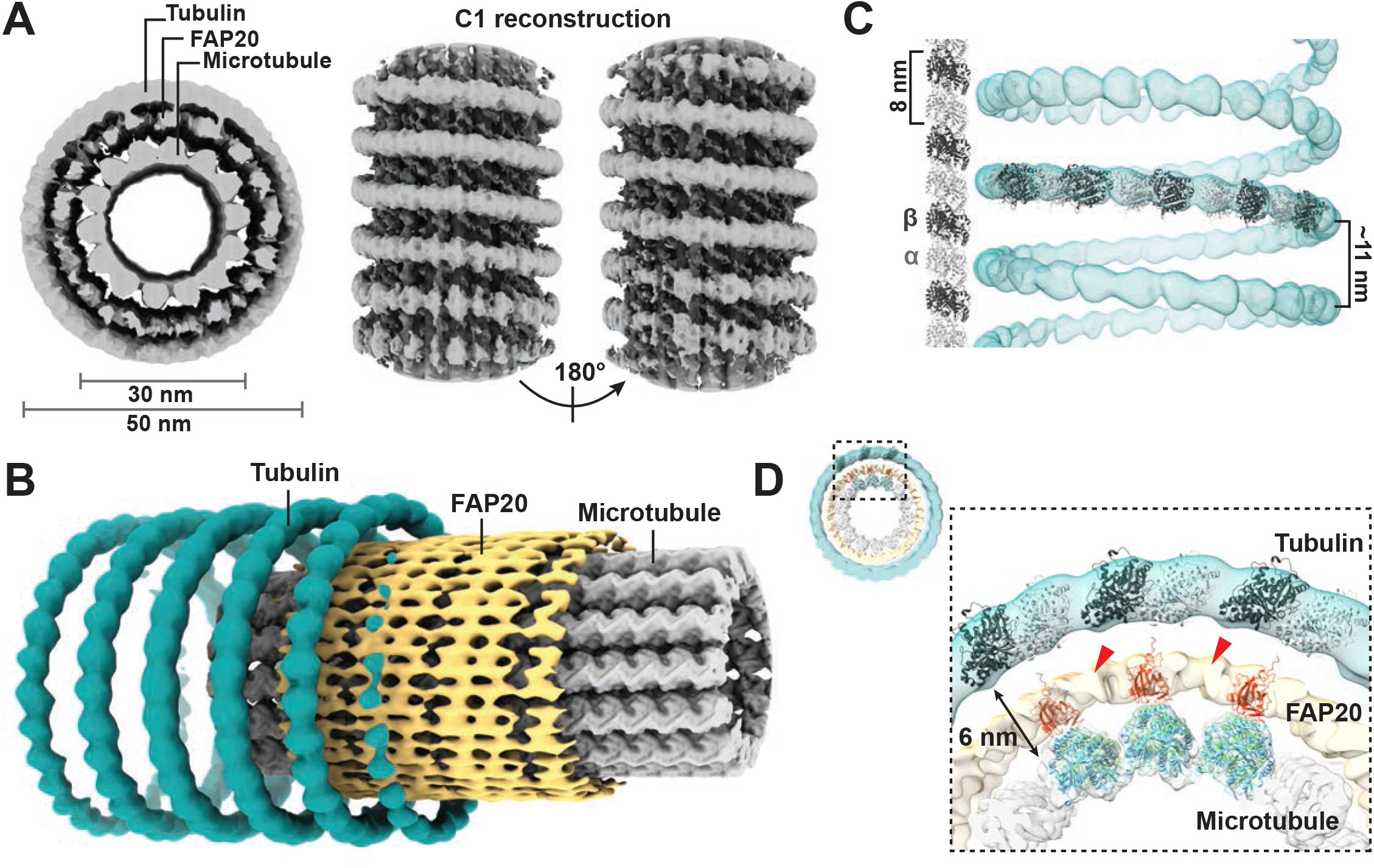
Cryo-electron microscopy maps and 3D reconstruction of microtubule: FAP20: tubulin complex. **(A)** Surface representation of C1 reconstruction of microtubule: FAP20: tubulin complex obtained in absence of GTP is shown in grey. Left: Cross section of the 3D reconstruction showing three concentric layers, representing the 14 protofilament microtubule (inner), FAP20 (middle) and tubulin dimers (outer). Right: Lateral views of the 3D reconstruction rotated by 180 degrees along the vertical axis of the microtubule to show the rings/spirals of FAP20: tubulin complex and absence of break in the rings/spirals corresponding to the microtubule seam. **(B)** Surface representation of symmetrized 3D reconstructions of each of the three concentric layers: microtubule (grey), FAP20 (yellow) and outer tubulin rings/spirals (teal) obtained by focused refinement after signal subtraction of the corresponding densities from particles. (See Methods section for more details). **(C)** Vertical versus horizontal protofilament arrangement model of tubulin dimers in the electron density map of outer tubulin layer from (B). Cartoon representations of atomic models of alpha and beta tubulin (obtained from AlphaFold2 prediction) are shown in light and dark grey respectively. Height of a tubulin dimer and approximate vertical distance of a tubulin spiral are indicated. **(D)** Cross section of symmetrized 3D reconstructions of microtubule, FAP20 and outer tubulin rings/spirals from (B) shown in the top left corner with a zoomed in version outlined in black. Electron density maps corresponding to three protofilaments of microtubule (inner) layer were fitted with three repeats of a model of FAP20 (red) bound to tubulin dimer (green/blue) obtained from AlphaFold2 prediction. Similarly, corresponding outer layer maps were fitted with three repeats of the tubulin dimer (light and dark grey) as shown in cartoon representation. The distance between the microtubule and outer tubulin layer is indicated with a double-sided arrow and the additional map densities corresponding to the FAP20 layer are marked with red arrows.

In summary, we have purified FAP20 and characterized its interaction with microtubules/tubulin dimers. Using biochemistry, TIRF and electron microscopy imaging we thus conclude that FAP20 mediates flexible interactions between microtubule and tubulin dimers possibly by undergoing conformational changes or oligomerisation. Our structure of microtubule:FAP20:tubulin complex determined in absence of GTP suggests that the binding of FAP20 to the microtubule lattice is specific.

## Discussion

The architecture of the doublet microtubule inner junction is fundamentally different from that of the outer junction and is chiefly composed of FAP20 and PACRG repeating units connecting the B-tubule to the A-tubule (Dymek *et al,* 2019). Previous studies on the inner junction have involved analysis of isolated doublet microtubules (Khalifa *et al,* 2020a; Ma *et al,* 2019), *Chlamydomonas* FAP20 and PACRG mutants (Dymek *et al,* 2019; Yanagisawa *et al,* 2014) and purified recombinant PACRG (Khan *et al,* 2021); however the properties of FAP20 have remained largely uncharacterized. In this study we recombinantly purified FAP20 and characterized its role in mediating the inner junction connections using TIRF and electron microscopy. Microtubule growth and dynamic assays in the presence of FAP20 revealed an increase in microtubule growth rate and length at high concentrations along with a stark reduction in frequency of observed catastrophe events. Similar stabilising lattice interactions are often observed in case of microtubule inner proteins (MIPs) associated with the doublet microtubules. *Chlamydomonas* mutants of MIPs, FAP45 and FAP52 show higher depolymerisation rates of the B-tubule compared to the wildtype (Owa *et al*, 2019). Turbidity studies with recombinant FAP85, a MIP associated with the A-tubule show reduced microtubule depolymerisation rates at lower temperatures (Kirima & Oiwa, 2018). Since FAP20 binds uniformly over the microtubule lattice *in vitro*, we conclude that this mode of microtubule polymer protection is unique to FAP20 among the microtubule doublet associated proteins that have been characterized so far.

In addition to microtubule stabilizing role of FAP20, our experiments revealed another salient property of FAP20, where it recruits free tubulin to the existing microtubule polymer. Recruitment of free tubulin to the existing polymer has been reported in microtubule repair mechanism and occurs without the aid of any MAPs (Schaedel *et al,* 2015). PACRG mediated tubulin recruitment has also been reported recently, however the tubulin has been shown to accumulate as distinct puncta (Khan *et al*, 2021). In contrast, our dynamic assays show that the FAP20 mediated tubulin recruitment is uniform and steadily increases over time in the presence of GTP (Figure 2A, 2C and Supplementary Movie 1). Through cryo-electron microscopy, we further show that FAP20 mediates flexible interactions with tubulin dimers in absence of GTP as well as larger assemblies of protofilaments, when GTP is added (Figure 3B and 3C). The higher order structure of microtubule:FAP20:tubulin obtained in presence of GTP appears similar to the ‘double walled microtubules’ nucleated in presence of polycations, peptides or histones (Erickson & Voter, 1976; Unger *et al,* 1990). The tubulin rings/spirals connected by FAP20 to the microtubule lattice obtained in absence of GTP resemble similar tubulin dimer decorations in the microtubule structures determined in complex with depolymerising kinesin-13 and Ska1 proteins (Benoit *et al,* 2018; Monda *et al,* 2017). The periodicity of FAP20 interaction with the microtubule lattice as evident by spacing between the FAP20-tubulin rings/spirals was found to be around 11 nm which is slightly higher than the 8 nm periodicity of FAP20 binding in the doublet microtubule (Dymek *et al*, 2019; Khalifa *et al*, 2020a; Ma *et al*, 2019). This difference can be attributed to a stagger of tubulin dimers in the outer ring as compared to the vertical axis of the microtubule due to FAP20 binding flexibly to alpha tubulin leading to different helical symmetries for the microtubule and outer ring (Figure 4C). In contrast, kinesin-13 binds between alpha and beta tubulin and the spirals formed of kinesin-13 and tubulin follow the helical path of the tubulin heterodimer in the microtubule lattice (Benoit *et al*, 2018; Tan *et al*, 2006).

While absence of FAP20 affects the beak structures in axonemes, it does not appear to be critical for other associated structures of the axoneme such as radial spokes, outer and inner dynein arms and the DRC (Yanagisawa *et al,* 2014). High resolution structures of the doublet microtubule suggest that FAP20 interacts with many inner junction proteins including PACRG, FAP52, FAP276, FAP106, FAP126 and FAP45 (Khalifa *et al,* 2020a; Ma *et al,* 2019). Extensive comparative analysis of expression patterns of these interacting proteins is lacking but studies point towards the relative independence of FAP20 expression. FAP52 mutants do not show decrease in levels of FAP20 or PACRG (Khalifa *et al*, 2020a) and reports have shown reduced levels of PACRG and Rib72 in FAP20 mutants (Yanagisawa *et al*, 2014), but not vice versa (Dymek *et al*, 2019) suggesting their recruitment to be sequential steps in a pathway. Since mutants of FAP20 or PACRG do not have incomplete B-tubules (Dymek *et al*, 2019), it is likely that both these proteins play a role in later stages of the doublet microtubule assembly after it has been initiated by the other proteins (Owa *et al*, 2019). However, absence of FAP20 in *Chlamydomonas* leads to flaying of isolated axonemes upon addition of ATP, indicating its importance in the stability of axonemes (Yanagisawa *et al,* 2014).

Based on the above inferences, we provide a basic model of assembly of the inner junction of doublet microtubules (Figure 5). As the B-tubule extends from the A-tubule anchored at the outer junction, several MIPs lining the B-tubule and ribbon of the doublet (FAP52, FAP276, FAP106, FAP126 and FAP45) act synergistically via tubulin post translational modifications and aid addition of requisite protofilaments for reaching the inner junction site (Ichikawa *et al*, 2017; Khalifa *et al*, 2020a; Ma *et al*, 2019). Mutations in these proteins are associated with ciliary defects in structure and function (Gegg *et al*, 2014; Jungnickel *et al*, 2018; Ta-Shma *et al*, 2015) emphasizing their role in cilia formation and maintenance. Upon expression, FAP20 and PACRG bind in different combinations to the partial doublet structure; it is still not clear if FAP20 and PACRG exist as or form heterodimers (Dymek *et al*, 2019; Khalifa *et al*, 2020a). However, at places along the inner junction where both of them bind, a stable intermediate is formed (Figure 5) which then propagates along the junction stabilising it. Although, we cannot rule out the role of a marker protein between (A13 and A1) such as FAP276 or tubulin PTMs at the carboxy-termini of B-tubules in recruiting FAP20 and PACRG to the position of the inner junction (Khalifa *et al*, 2020a) our *in vitro* reconstitution studies as well as AlphaFold2 models suggest that FAP20 is a key molecule in mediating the interaction between the protofilaments of A- and B-tubule and PACRG. Although, higher resolution structures of *in vitro* reconstituted complexes at different stages of doublet assembly will be required to understand and validate the assembly mechanism in detail, the purification and reconstitution experiments with FAP20 in this study lay the foundation and provide a step closer towards polymerizing doublet microtubules *in vitro*.

**Figure 5.**
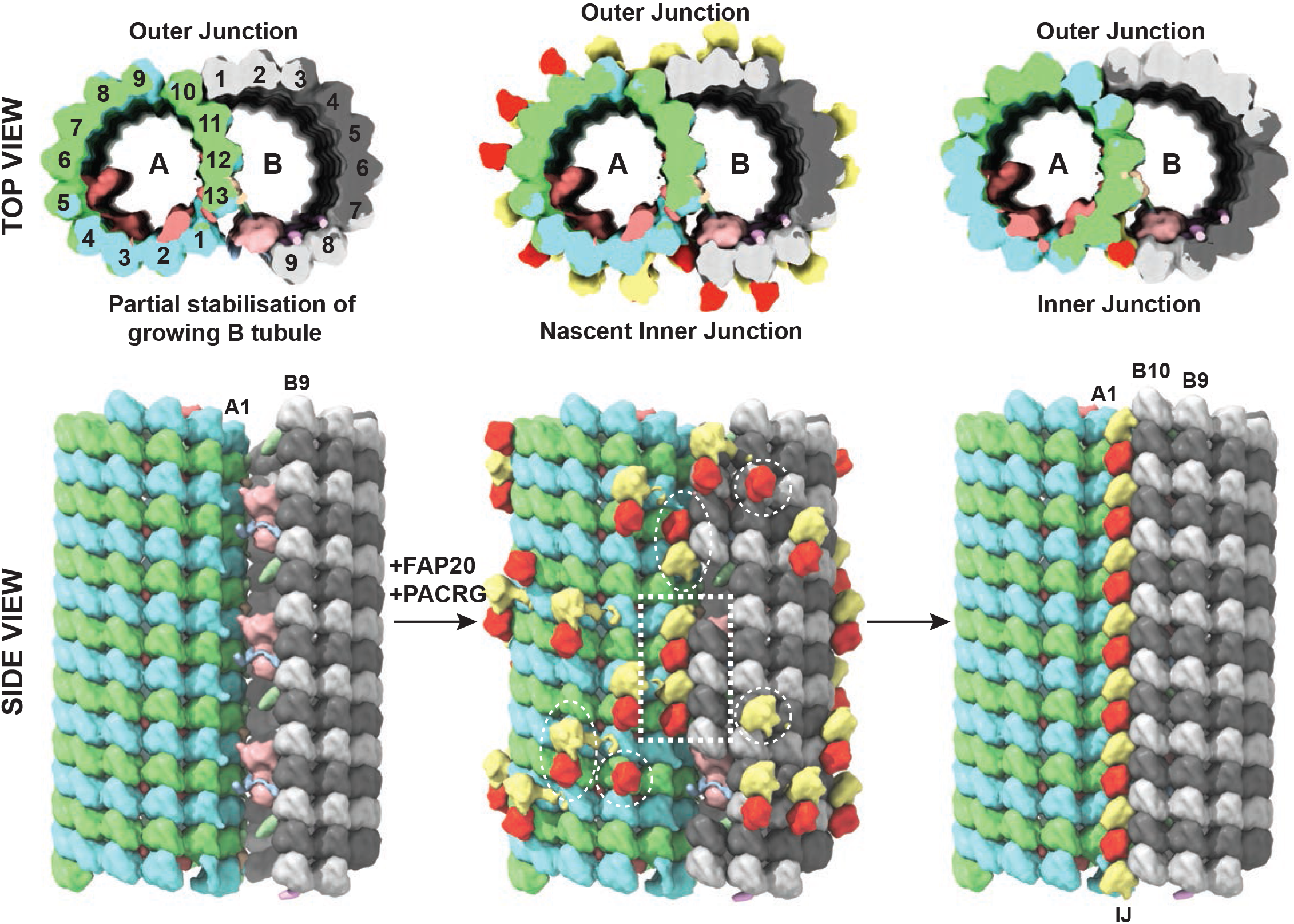
Model for assembly of inner junction and closure of doublet microtubule. Schematic model illustrating the steps of formation of the inner junction. Cross sections are shown on the top with the corresponding lateral views depicted below. Partial stabilisation of B-tubule with proteins implicated in vital contacts at the inner junction (left) followed by stochastic formation of a nascent inner junction in the presence of FAP20 and PACRG along the joining points of B- and A-tubule (middle) and finally the propagation of this stable inner junction segment along the longitudinal axis leading to the formation of the complete inner junction and the closure of B-tubule of the doublet microtubule (right). Density maps generated using the doublet microtubule model (PDB: 6u42) in Chimera are depicted as surface representation with alpha and beta tubulins from the A-tubule shown in green and blue respectively, FAP20 and PACRG in red and yellow respectively and alpha and beta tubulin from the B-tubule in light and dark grey respectively. The proteins of the inner junction portrayed here include FAP45 (pink), FAP52 (peach), FAP106 (light green), FAP126 (beige), FAP276 (light blue) and RIB72 (salmon). The different possible combinations of FAP20 and PACRG interactions with the microtubule lattice are shown in white dashed circles/ovals, while the stable inner junction segment is outlined in a white dashed rectangle.

## Materials and Methods

### Purification of proteins

Tubulin was purified by a series of polymerization and depolymerization steps from goat brain using a standard protocol (Castoldi & Popov, 2003). The coding sequence for FAP20 was amplified from *C. reinhardtii* cDNA and cloned into pET28A vector between *BamH1* and *Nde1* restriction sites with a carboxy terminal deca-histidine tag. The FAP20 pET28A vector was transformed into Rosetta *E. coli* cells and a single transformed colony was used for inoculation in 25 ml of Luria Bertani broth for overnight growth as primary inoculum. 25 ml of this primary inoculum was added to 1L of Terrific Broth medium and incubated at 37°C for 4-6 hours. 0.5 mM IPTG was added for induction of 10X His tagged FAP20 and the culture was left: under shaking conditions at 150 rpm, 25°C overnight. Next day, the cells were harvested and resuspended in buffer containing 50mM HEPES pH 7.5, 500mM KCl, 10 mM imidazole, 5mM β mercaptoethanol, 100mM PMSF and EDTA free Protease Inhibitor Cocktail tablets (Roche, Cat No.11836170001) using a manual dounce. The resuspended cells were lysed by sonication followed by centrifugation to remove cell debris. The clarified lysate was loaded at a slow flow rate onto an equilibrated Ni-NTA column. The loaded column was washed with 5 column volumes of wash buffer containing 50mM HEPES pH 7.5, 250 mM KCl, 40 mM imidazole, 1 mM ATP, 2 mM MgCl2 and 5 mM β mercaptoethanol to remove contaminating proteins. Column bound proteins were then eluted using a gradient of 40 mM to 350 mM imidazole in 50mM HEPES pH 7.5, 5mM β mercaptoethanol buffer. The protein that eluted at a concentration of around 200mM imidazole corresponded to FAP20 and was passed through pre-equilibrated Q column to remove other contaminants. We observed that FAP20 did not bind to Q column, however the flowthrough was devoid of other contaminating proteins. The Q column flowthrough and wash was then passed through a Ni-NTA column again and eluted in a smaller volume, which was then concentrated using a centricon with 10 kDa molecular weight cut off. Imidazole was removed by 2 rounds of dilution and concentration with buffer containing no imidazole. The purity of the protein was confirmed by SDS page which showed a single band at around 20 kDa. Small aliquots (5-10 μL) of proteins were snap frozen in liquid nitrogen and stored at - 80°C

### Microtubule co-pelleting assay

Purified goat brain tubulin (20 μM) was polymerized in BRB80 buffer (80mM PIPES, 1mM MgCl_2_, 1mM EGTA, pH 6.8 set with KOH) with 2 mM GTP followed by addition of 20 μM taxol after 30 minutes and incubated at 37°C for 2-3 hours. Similarly, microtubules were also polymerized with a slowly hydrolysable GTP homologue, 0.4 mM GMPCPP but without taxol. Microtubules supplemented with 2mM GTPγS were incubated at 37°C overnight in BRB80 buffer. 5μM FAP20 was incubated with polymerized microtubules for 20 minutes at room temperature followed by centrifugation in a TLA100 rotor at 279,000 x g for 10min at 35°C layered over a 60% glycerol cushion (60mL 100% glycerol, 20mL dd-water, 20mL 5X BRB80). The pellet was dissolved in 40uL of warm BRB80 and was analyzed along with the supernatant on a 10% SDS-PAGE. 10 μM kinesin-1 (1-324 G234A) was incubated with microtubules for 20 minutes at room temperature and pelleted down following the same protocol. The comparative densitometric analysis of FAP20, kinesin and tubulin were done using ImageJ software.

### *In vitro* reconstitution

1.5 mm coverslips were cleaned by sonication in acetone and stored in 80 % ethanol. Short GMPCPP microtubule seeds were prepared by polymerization at 37°C for 30 minutes, spun on warm 50% BRB80 sucrose cushion and resuspended in warm (37°C) BRB80 buffer. These seeds were attached by means of BSA-biotin-streptavidin linkage onto the coverslips in a flow chamber. Assay mix consisting of BRB80 buffer supplemented with β-casein (62μg/ml), 10mM β-mercaptoethanol, 2mM GTP, 0.1% methyl cellulose and oxygen scavenging system was prepared in ice. To a 20μl aliquot of this mix, 10uM tubulin (spiked with 647-Alexa labelled tubulin) and FAP20 was added and centrifuged to remove aggregates. The range of concentrations of FAP20 (0.05μM to 5μM) used was based on the stoichiometric ratio of bound FAP20 to tubulin (0.25:1) obtained from microtubule copelleting experiments (Supplementary Figure 1C). The supernatant was warmed to 25°C and flowed into the flow chamber and immediately visualized using a Nikon TIRF-2 system at 25°C. Image sequences were collected at 561 and 647nm excitation wavelengths with an exposure time of 100 milliseconds at an interval of 2 seconds for 10 minutes.

### Data Analysis

The TIRF imaging data were analyzed with Fiji software including drift correction when necessary, using StackReg (Thevenaz *et al,* 1998) rigid body transformation. Microtubule growth rates, growth lengths and catastrophe frequencies were calculated from kymographs plotted from single isolated microtubules manually selected in the maximum intensity projections of the movie frames. Growth length refers to the total length of the newly polymerized segment of the microtubule for the duration of the experiment. In case of higher concentrations of FAP20 (2.5 and 5μM) catastrophe events were considered when microtubules shortened by at least a third of their growth length. Intensity measurements were made in Fiji over time or microtubule length for manually selected single isolated microtubules. Intensity was normalized by subtracting intensity values over the same selected region between microtubules and an adjacent area in the field of view. For each condition, data were analyzed from 3 independent experiments and graphs were plotted using GraphPad Prism version 9. SuperPlots were generated as described in (Lord *et al*, 2020)

### Sample preparation and data collection for cryo-electron microscopy

Quantifoil gold holey carbon grids R 1.2/1.3 were glow discharged in a GloQube Glow Discharge (Quorum Technologies) system for 90 seconds. Sample application and vitrification was carried out in a Vitrobot Mark IV (ThermoFisher Scientific) set to 25°C and 100% humidity using a Whatmann No. 1 filter paper for blotting.

50μM tubulin was polymerized with 1mM GMPCPP at 37°C for 2 hours, spun on warm 50% BRB80 sucrose cushion and resuspended in warm BRB80 buffer. A mix of 10μM tubulin and 10μM FAP20 in BRB80 buffer was prepared in ice and centrifuged at 4°C to remove aggregates. The supernatant was kept in ice and warmed only prior to application on the grids. 1μL of the GMPCPP microtubule seeds was applied on the grid mounted in the Vitrobot. After a brief incubation of 10-20 seconds to allow for adsorption of the seeds onto the grid, 3μL of pre-warmed FAP20-tubulin mix was applied to the same grid. The grids were then blotted after an incubation time of 1 minute with a blot force of 0 and blot time of 3 seconds before plunging into liquid ethane. The frozen grids were transferred to an Autogrid under liquid nitrogen conditions and mounted into a Titan Krios (FEI) operating at 300keV. Data collection was carried out using the automated pipeline of EPU software system with the Falcon III detector in integration mode at the National CryoEM facility, Bangalore. Images were collected at a magnification of 47,000x resulting in a pixel size of 1.78Å and defocus range between −2 to −3.2 μm. Each image had a total electron flux of 62.15 electrons/pixel/sec resulting in a total dose of 39.55 electrons/Å^2^ and 20 movie frames were stored.

### Cryo-EM image processing

Summed images were manually inspected for presence of contaminating ice and filament quality and poor-quality micrographs were discarded. Motion correction of movie frames was carried out using Unblur (Grant & Grigorieff, 2015) or MotionCor2 (Zheng *et al,* 2017) and CTF estimation was done on dose-weighted motion corrected summed images using GCTF (Zhang, 2016). All subsequent image processing steps were carried out in Relion v3.1 (Scheres, 2012a, 2012b). Helical segments were manually picked using Relion’s helical picker (He & Scheres, 2017) and 2x binned particles were extracted using a box size of ~780Å and overlapping inter-box distance of 82 Å. Microtubule segments containing different protofilament numbers were segregated using supervised 3D classification involving comparison with references representing microtubules made of 11 to 16 protofilaments. The 14 protofilament microtubules formed the major fraction of the population and were selected for further processing. In case of microtubule:FAP20:tubulin with GTP dataset, this subset of unbinned 14 protofilament containing microtubule segments was used for several runs of 3D refinement using a combination of cylinder and low-pass filtered microtubule as references, with and without imposing helical symmetry and in absence and presence of soft masks but a well resolved consensus structure could not be obtained (Supplementary Figure 4B). This could be because of high levels of uncontrolled heterogeneity in the sample that could not be segregated by 3D classification. For the microtubule:FAP20:tubulin (without GTP) dataset, the set of unbinned 14 protofilament microtubules was used for 3D refinement with helical symmetry using a cylinder with inner and outer diameters of 18 and 55 nm respectively as reference. The obtained model was again refined with imposed helical symmetry and a soft mask extending between 35 to 60 nm and containing 30% of the filament to improve the alignment of the outer tubulin rings. This model was then used for 3D refinement without symmetry starting with a sampling rate of 1.8 degrees to give the C1 reconstruction (Figure 4A). Examination of the obtained structure revealed that the different layers of microtubule, FAP20 and outer tubulin rings possibly followed different helical parameters leading to improved features in parts of the reconstruction depending on the focus and masks applied. In order to improve reconstruction of each layer, signal subtraction was applied to the particles using customised masks focussing on each concentric layer. Refinement of subtracted particles containing only the microtubule was carried out with helical symmetry using GMPCPP microtubule (EMDB 7973) low pass filtered to 15Å as the reference. A map sharpening B-factor of −163 Å^2^ was applied to the final map. For subtracted particles focused on FAP20 and outer tubulin ring layers, 3D classification with and without symmetry was attempted to solve the problem of heterogeneity but was unsuccessful. The subtracted particles for each layer (FAP20 and outer tubulin helix) were then independently refined with helical symmetry using suitable references and masks (Supplementary Figure 5A). The data collection and refinement statistics are provided in Table 1. The electron density map for C1 reconstruction of microtubule:FAP20:tubulin complex without GTP is deposited in the Electron Microscopy Data bank with deposition number EMD-34836.

**Table 1:**
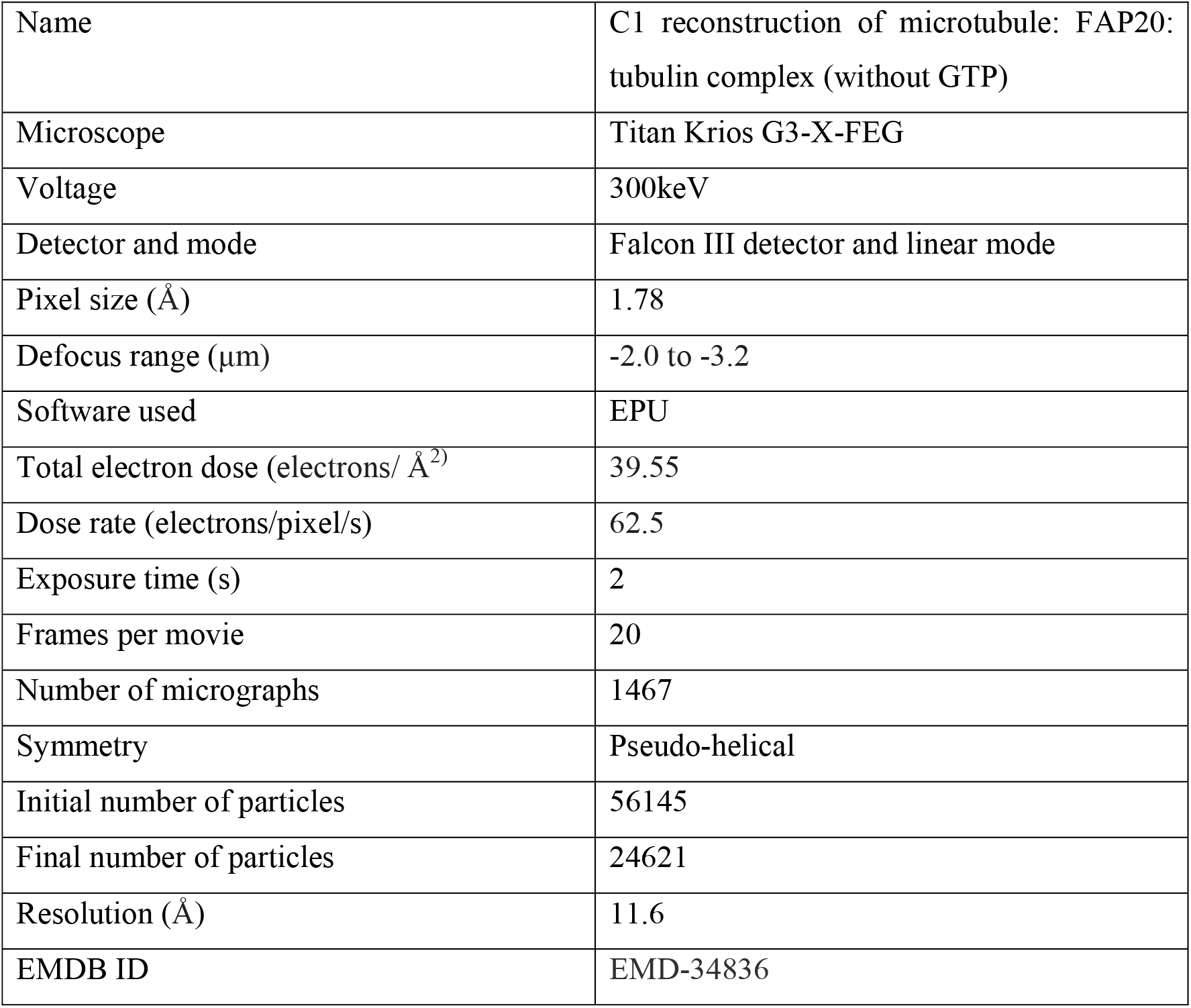
Data collection and refinement statistics.

|  |  |
| --- | --- |
| Name | C1 reconstruction of microtubule: FAP20: tubulin complex (without GTP) |
| Microscope | Titan Krios G3-X-FEG |
| Voltage | 300keV |
| Detector and mode | Falcon III detector and linear mode |
| Pixel size (Å) | 1.78 |
| Defocus range (μm) | -2.0 to -3.2 |
| Software used | EPU |
| Total electron dose (electrons/ Å<sup>2</sup>) | 39.55 |
| Dose rate (electrons/pixel/s) | 62.5 |
| Exposure time (s) | 2 |
| Frames per movie | 20 |
| Number of micrographs | 1467 |
| Symmetry | Pseudo-helical |
| Initial number of particles | 56145 |
| Final number of particles | 24621 |
| Resolution (Å) | 11.6 |
| EMDB ID | EMD-34836 |

Since the resolution of the maps obtained were not sufficiently high enough to build atomic models unambiguously, AlphaFold2 was used to obtain atomic models that could be fitted into the electron maps. The models obtained were docked into the map using Chimera’s ‘Fit in Map’ option. Superposition of atomic models were carried out in Coot (Emsley *et al,* 2010) and images were made using Chimera (Goddard *et al,* 2007; Pettersen *et al,* 2004) and ChimeraX (Goddard *et al,* 2018; Pettersen *et al,* 2021)

### AlphaFold2 prediction and analysis

Protein sequences for goat brain tubulin (Uniprot IDs A0A452F3L8 for alpha tubulin and A0A452FIJ2 for beta tubulin) and FAP20 (Uniprot ID A8IU92) were used as inputs for locally installed AlphaFold2 (Evans *et al*, 2021; Jumper *et al*, 2021b). The model with the highest confidence calculated as a combination of pTM (predicted TM-score) and ipTM (interface predicted TM-score) was used for fitting and analyses.

## Acknowledgements

The authors acknowledge the National Cryo-EM facility at inStem, Bangalore, the funding by the B-life grant from the Department of Biotechnology (DBT/PR12422/MED/31/287/2014) and K.R. Vinothkumar (National Centre for Biological Sciences, India) for the help in EM data collection. M.B. would like to thank David Houldershaw and Birkbeck IT facility for computational support. We thank K.R. Vinothkumar, Sudarshan Gadadhar and Aakash Mukhopadhyay for comments on the manuscript. M.B acknowledges the SERB-NPDF fellowship (PDF/2016/002682). M.S acknowledges funding support from inStem core grants from the Department of Biotechnology, India, DBT/Wellcome Trust India Alliance Intermediate Fellowship (IA/I/14/2/501533), EMBO Young Investigator Programme award, CEFIPRA (5703-1) from the Department of Science and Technology, SERB-EMR grant (CRG/2019/003246) and DBT-BIRAC (BT/PR40389/COT/142/6/2020) grant.

## Author contributions

M.B and M.S conceived the project. M.B purified FAP20, performed TIRF and cryoEM work, analysed the data and wrote the paper. S.P established the protocol for purification of FAP20. A.D performed the co-sedimentation assays described in Supplementary Figure 1. M.S acquired the funding, supervised the work and wrote the paper.

## Supplementary Figures

**Supplementary Figure 1. FAP20 purification and binding with microtubules.**

**(A)** Coomassie stained SDS-PAGE gel showing bands corresponding to the purified FAP20 and different concentrations of BSA standard used in estimation of protein concentration.

**(B)** Coomassie stained SDS-PAGE gel showing microtubule co-pelleting experiments with FAP20 and Kinesin-1 (K349-G234A). Inp = Input; Sup = Supernatant; P = Pellet.

**(C)** Quantification of microtubule co-pelleting experiments showing the fraction of FAP20 or Kinesin-1 bound to the tubulin. Mean values are shown in the form of bars with the error bars representing standard error of mean calculated from three independent experiments.

**Supplementary Figure 2. Effect of FAP20 binding on microtubule dynamics.**

**(A) and (B)** Quantification of growth rates and growth lengths of microtubules upon binding of different concentrations of FAP20 (0 to 5μM) for each experiment, as indicated. Data is plotted as individual values in the form of coloured circles with mean and error bars representing standard error of mean. (Number of growth events analysed in each experiment; N1 = 50, N2 = 31, N3 = 54: 0μM FAP20; N1=35, N2=41, N3=54: 0.05μM FAP20; N1=38,N2=33,N3=66: 0.25 μM FAP20; N1=31, N2=27, N3=60: 0.5μM FAP20; N1=30, N2=58, N3=76: 2.5μM FAP20; N1=36, N2=39, N3=75: 5μM FAP20. Number of microtubules analysed for growth length in each experiment N1=32, N2=23, N3=42: 0μM FAP20; N1=30, N2=27, N3=44: 0.05μM FAP20; N1=28, N2=27, N3=53: 0.25 μM FAP20; N1=28, N2=21, N3=45: 0.5μM FAP20; N1=27, N2=56, N3=42: 2.5μM FAP20; N1=31, N2=37, N3=42: 5μM FAP20. *p<0.05, **p<0.01, ***p<0.001, and ****p<0.0001, Kruskal-Wallis test followed by Dunn’s multiple comparisons test.)

**(C)** Bar graph showing number of microtubules from each experiment undergoing different catastrophe frequencies at the indicated FAP20 concentration (N1 = 32, N2= 23, N3=42 : 0μM FAP20, N1=30, N2=27, N3=44 : 0.05μM FAP20, N1=28, N2=27, N3=53: 0.25 μM FAP20, N1=28, N2=21, N3=45: 0.5μM FAP20, N1=27,N2=56, N3=42 : 2.5μM FAP20, N1=31,N2=37, N3=42: 5μM FAP20).

**Supplementary Figure 3. Recruitment patterns of cy5 labelled tubulin by 5μM FAP20 on the microtubule lattice.**

**(A)** Kymograph showing puncta of cy5-labelled tubulin along the GMPCPP stabilised seed as well as the growing microtubule formed in presence of GTP, indicated by white arrows.

**(B)** Kymograph showing uniform recruitment of cy5-labelled tubulin along the GMPCPP stabilised seed in presence of GTP, with the starting point as indicated by a white arrow.

**(C)** Kymograph showing higher levels of recruitment along the growing microtubule as compared to the GMPCPP stabilised seed in presence of GTP.

**(D) and (E)** Representative kymographs showing uniform recruitment along the GMPCPP seed in absence of GTP.

In each case, ‘seed’ channel represents excitation at 561 nm and ‘tubulin’ channel represents excitation at 641nm. Dotted white lines in the ‘tubulin’ channel denote the ends of GMPCPP stabilised seeds. Scale bars represent 5 μm along X axis and 2 minutes along the Y axis.

**Supplementary Figure 4. Electron microscopy-based analysis of microtubule: FAP20: tubulin complex.**

**(A)** 2D power spectrum of GMPCPP microtubule segments (left), in presence of FAP20 and tubulin with GTP (centre) and without GTP (right). All three spectra show strong layer lines at orders of 1/4 nm-1 (black arrows) indicating similar arrangement of tubulin monomers along the microtubule lattice. The spectrum of microtubule with FAP20 and tubulin in absence of GTP (right) shows an additional layer line corresponding to 1/10 nm-1 (red arrows) suggesting that the FAP20-tubulin complex follows a different helical pattern unlike the tubulin monomer in the microtubule.

**(B)** Data processing pipeline to obtain a 3D reconstruction (grey) of FAP20-tubulin bound microtubule complex in presence of GTP (See Methods section for details). Inset: Cross-section of the 3D reconstruction of the structure obtained without imposing symmetry (C1 refinement) shown in grey superimposed with a 10Å low pass filtered 14 protofilament microtubule map (EMD 7973) shown in cyan.

**Supplementary Figure 5. Electron microscopy and Alphafold2 prediction-based analysis of FAP20-tubulin-microtubule complex in absence of GTP.**

**(A)** Schematic outline of the processing pipeline for focused refinement of each concentric layer of the complex (See Methods section for details). 3D classification attempts to identify subpopulations of particles showing different arrangements focussed on outer tubulin and FAP20 layers from subtracted segments are shown in blue and pink boxes respectively.

**(B)** Fourier shell correlation plots of the half maps for C1 reconstruction (Figure 4A) of the FAP20-tubulin-microtubule complex in absence of GTP and symmetrized reconstructions of each of the layers of microtubule, FAP20 and outer tubulin (Figure 4B) obtained after focused refinements of the corresponding layer from signal subtracted particles. The resolutions were estimated based on 0.143 criterion of half maps.

**(C)** Superposition of the top atomic model based on confidence assessed by ipTM+pTM score (See Methods for details) obtained from AlphaFold2 for FAP20 + tubulin dimer shown in red upon the unit of the inner junction (Chains G, H, I of Bundle 1: A-tubule; b of Bundle 12: FAP20; c of Bundle 12:PACRG; L, M, N of Bundle 12: B-tubule from PDB ID: 6u42) shown in grey. The FAP20 chain was used for superposition.

**(D)** The top 5 models obtained from AlphaFold2 for FAP20+tubulin dimer are shown with alpha tubulin chains used for superposition on the top model. The corresponding confidence scores are shown in the associated table.

**Supplementary Movies**

**Supplementary Movie 1: Stabilisation of the growing microtubule and recruitment of free tubulin to the microtubule lattice by FAP20 in presence of GTP.**

Dynamic microtubules (cyan) grown from GMPCPP stabilised cy3-labelled seeds (magenta) and 10μM cy5-labelled tubulin (top) and in presence of 5μM unlabelled FAP20 (bottom) with GTP. Corresponding movie frames from the cy5-tubulin channel only are shown on the right. Time is in min:sec, Playback: 50fps

**Supplementary Movie 2: FAP20 recruits free tubulin to the microtubule lattice in the absence of GTP.** GMPCPP stabilised cy3-labelled seeds (magenta) incubated with 10μM cy5-labelled tubulin only (top) and with 5μM unlabelled FAP20 (bottom) in absence of GTP. Corresponding movie frames from the cy5-tubulin channel only are shown on the right. Time is in min:sec, Playback: 50fps

